# A Wildlife Health Outbreak Response Table-top Exercise for Pandemic Preparedness Planning

**DOI:** 10.1101/2025.04.01.646690

**Authors:** Vida Ahyong, Patrick Ayscue, Rebecca Gowen, Natasha Spottiswoode, Charles E. Alex, Eman Anis, Daniel P. Beiting, Michelle Gibison, Eva I. Gordián-Rivera, Sabrina S. Greening, Lucy Keatts, R. Scott Larsen, Erica Miller, Brian Niewinski, Roderick B. Gagne, Cristina M. Tato, Julie Ellis, Lisa Murphy

## Abstract

Zoonotic diseases have received significantly more attention over the last few decades, emerging with increasing frequency and causing the majority of notable disease outbreaks in this century, including the COVID-19 pandemic. As human activities and shifting climate patterns induce changes in the environment that alter habitat and range of reservoir species, the potential for human and animal interactions will increase and enhance the opportunity for spillover. Thus, any emergency response preparedness planning must take into account the function and coordination of agencies across the sectors of human, animal and environmental health. Within the Commonwealth of Pennsylvania a table-top exercise was performed to evaluate a multi-agency response during a hypothetical zoonotic disease investigation. The exercise was evaluated by the participants to gain feedback on the overall process and lessons learned. Here, we describe the tabletop exercise scenario and the insights gained. We found that differences in operational structure create challenges for interdepartmental communication and in the ability to resource a coordinated response, highlighting opportunities to develop infrastructure that will facilitate future actions. A set of recommendations are outlined that may enhance cross-agency activities and promote more effective and efficient emergency response.

## Background

Of the myriad of known human pathogens more than 60% are transmitted from animals (Rahman et al., 2020). Zoonotic diseases have increased significantly in recent decades including notable examples like Middle East respiratory syndrome, Crimean-Congo hemorrhagic fever, and filoviruses such as Ebola and Marburg. Additionally, the global pandemic caused by SARS CoV-2 and the ongoing outbreaks of highly pathogenic avian influenza virus (HPAI) underscore the escalating threat posed by these pathogens (Ison & Marrazzo, 2025; Koopmans et al., 2024; Skowron et al., 2023). Zoonotic diseases cause an estimated 2.5 billion human cases of illness globally and 2.7 million deaths annually (Van Der Westhuizen et al., 2023) and can cause massive social and economic disruptions (McKibbin & Fernando, 2023), such as those observed during the COVID-19 pandemic and the products stemming from outbreaks of HPAI.

Human activities, including deforestation, trade in animal products, agriculture, and globalization, have accelerated the emergence, transmission, and proliferation of zoonotic pathogens by intensifying interactions between humans and animals (Allen et al., 2017; Weiss & McMichael, 2004). Furthermore, climate change-induced shifts in average temperatures and precipitation patterns are altering the habitats and ranges of reservoir species and pathogen vectors, exacerbating the spread of infectious diseases (Altizer et al., 2013). The complex set of drivers across systems means that early identification and response to emerging zoonotic diseases requires coordination between multiple experts in human, animal, and environmental health (Aarestrup et al., 2021; Sharan et al., 2023). Often these sectors are managed by independent agencies with focused mandates and without established pathways for communication and data sharing between them. For example, in the United States wildlife health, domestic animal health and public health are managed by separate agencies, primarily at the state or local level. We learned from SARS-CoV-2 the importance of effective responses at the local municipality and state levels, and the need for coordination across entities. Thus, bringing together representatives from multiple agencies to evaluate response to a zoonotic disease is a critical step in future mitigation of emerging diseases.

A strategy for evaluating multi-agency response to a zoonotic disease is to bring together relevant stakeholders to conduct table-top exercises that run through a prepared simulated emerging disease event (Chugh et al., 2024). Pennsylvania is an important location for such an exercise as it is located in the Northeast United States, a known hotspot for emerging infectious diseases (Fauci, 2005; Jones et al., 2008). Therefore we set out to conduct a table-top exercise around a zoonotic disease outbreak within a local municipality in Pennsylvania. In collaboration with the University of Pennsylvania’s School of Veterinary Medicine, we brought together an interdisciplinary group of human and animal health professionals, including the Pennsylvania Department of Public Health, the Pennsylvania Game Commission, privately run wildlife rehabilitation groups and University of Pennsylvania’s Wildlife Futures program. An exercise was designed around a wildlife disease event and implemented in September 2023 as an in-person round-table exercise.

## Description

The aim of the exercise was to understand how we currently function as a response network during a developing, potentially zoonotic wildlife disease event in the Commonwealth of Pennsylvania. Two main objectives were defined to focus the exercise and to help understand current gaps and challenges that may prevent effective emergency response. These objectives were to 1) identify areas where collaboration can be fostered around surveillance and response, and 2) identify operational gaps where additional planning is needed across groups in risk communication, surge capacity, and outbreak management. The exercise was hosted at the Pennsylvania Fish & Boat Commission headquarters in Harrisburg, PA on September 19, 2023. Participants included representatives from both public and private organizations that encompassed human and animal health missions, and included the University of Pennsylvania School of Veterinary Medicine’s Wildlife Futures Program and Pennsylvania Animal Diagnostic Laboratory System (PADLS), Pennsylvania Game Commission, Pennsylvania Department of Health, Pennsylvania Fish and Boat Commission, Pennsylvania Department of Agriculture, Wildlife Conservation Society, University of California San Francisco Division of Infectious Diseases and Chan Zuckerberg Biohub San Francisco.

### Scenario and process

The exercise was conducted with prompts progressively revealed to the participants with discussion after each reveal. As each prompt was presented to the group, participants were asked to evaluate the situation, decide on any action(s) to be taken, and identify any additional information needed. Discussion time was monitored to ensure completion of the exercise, along with debriefing and formulation of next steps, within a 6-hour timeframe.

#### Prompt 1

In the context of several endemic vector-borne diseases, wildlife surveillance indicates a greater than typical number of dead deer, especially in proximity to waterways, occurring in the greater Northeast (NE) region of PA.

#### Prompt 2

Initial exam during deer necropsy and additional diagnostic tests indicate a possible infectious cause that remains unidentified.

#### Prompt 3

A death of a 63 year old male human is reported from a suspected case of a tick-borne illness of unknown etiology. He initially presented to his primary care clinician with fever, diarrhea, and myalgia on July 1 following a camping trip with family on private property in NE PA the previous week. History indicated that he and other family members had contact with multiple deer carcasses encountered while camping in June.

#### Prompt 4

An infectious disease physician reports she is consulting on three additional cases with similar symptomology and unknown etiology in the hospital, all with histories of outdoor exposure in NE PA in the preceding weeks.

#### Prompt 5

Pathogen metagenomic sequencing results from deer and humans recovered reads aligning to a Bandavirus genome in the Bunyavirales order in all of the cases and none of the controls. This is a tick-borne virus with zoonotic potential and the ability to spread person-to-person through contact with infectious bodily fluids.

The group discussed the prompts with facilitators probing relevant areas for in-depth discussion with questions. Participants were encouraged to draw from experiences related to past events in which they were involved. Following completion of the exercise, the group was asked to evaluate the effectiveness of the response across partners, and to identify operational gaps.

### Evaluation survey

In the week following the exercise, a survey was distributed across all the participants with the aim of assessing the overall utility of the exercise and informing design of similar exercises in the future. The survey consisted of 16 questions including a mixture of multiple-choice, Likert scale and free text responses. A copy of the survey is provided in Appendix 1.

## Results

Discussion during the exercise revealed strengths, challenges and opportunities for improving intra-agency and multisectoral response to zoonotic infectious disease outbreaks.

### How effective was coordinated surveillance among partners?

Agencies have tended to work in awareness of each other’s activities, but independently of each other with limited coordination. Each organization operates within their own standard operating procedures (SOPs) for an infectious disease event, which are driven by individual resources, manpower, local regulations, and agency remit and priorities. This was highlighted by one participant who noted “We are often handcuffed by regulatory limitations when it comes to sharing personnel or resources across agencies.” Informal lines of communication do exist between agencies, however there is no mandate to disseminate information nor is there a designated information officer among them to gather intelligence for messaging and decision making. Participants specifically stated in the exercise that the decision to share information across agencies would be made by individuals and that the first line of contact would likely be a call to a known colleague at another agency. There is no infrastructure or database shared across state agencies to coordinate response outside of an emergency operations center, including for increased infectious disease surveillance efforts.

### How effectively were resource needs identified?

Participants identified required resources for the response scenario including access to material resources, clarity on regulatory authority, effective public-facing and interagency communications, and additional personnel resources. However, coordination of testing, response, or collaboration with agencies with complementary expertise had no existing SOPs. Local county authorities have oversight of funding and expenditures for individual agencies, and even bureaus within a state-run organization may have independent budgeting structures, making it difficult to quickly leverage existing monies or personnel for emerging priorities. There are no formal mechanisms in place at the agency level for mobilizing resources in an emergency for cross-agency collaboration and support. Limited standardized processes are in place for working with federal authorities for key public health interventions, such as determining quarantine mandates for novel zoonotic events.

### Were there barriers to sharing data across organizations?

While some agencies have a robust reporting system that would be of value to catalyze earlier activation of both public and private partners for an emergency response, the data are often protected under the authority of local municipalities or privacy laws, and are therefore not easily accessed, released or shared. Partner agencies were often unfamiliar with the types or format of relevant data collected by the other agencies. Agencies are not incentivized to share information, as it may jeopardize their funding or go against local mandates. These issues extend from local into regional interstate agency coordination during a wildlife event. Additionally, it was noted that there is a lack of formal guidelines for joint agency public messaging when a wildlife event has broader implications for public health and safety.

### Evaluation survey

All ten participants who completed the survey found the exercise to be beneficial with five ranking it a four out of five or “Very Useful” and five ranking it a five out of five or “Extremely Useful” (Figure 1A and B). A response by one participant highlighted that “The workshop was fantastic for facilitating interesting and useful discussions about various stakeholders’ roles in a wildlife/zoonotic disease outbreak, and identifying opportunities for clearer communication and cooperation among them.” All survey respondents were positive about participating in future exercises with nine out of ten respondents ranking a five out of five likelihood of participating in future similar exercises and a single participant indicating a four out of five (Figure 1C). Ideas for improving the exercise included “real time graphics” or other approaches to bringing together the complete picture of the event and identifying gaps. Multiple participants indicated more time should be given to future exercises.

**Figure 1.**
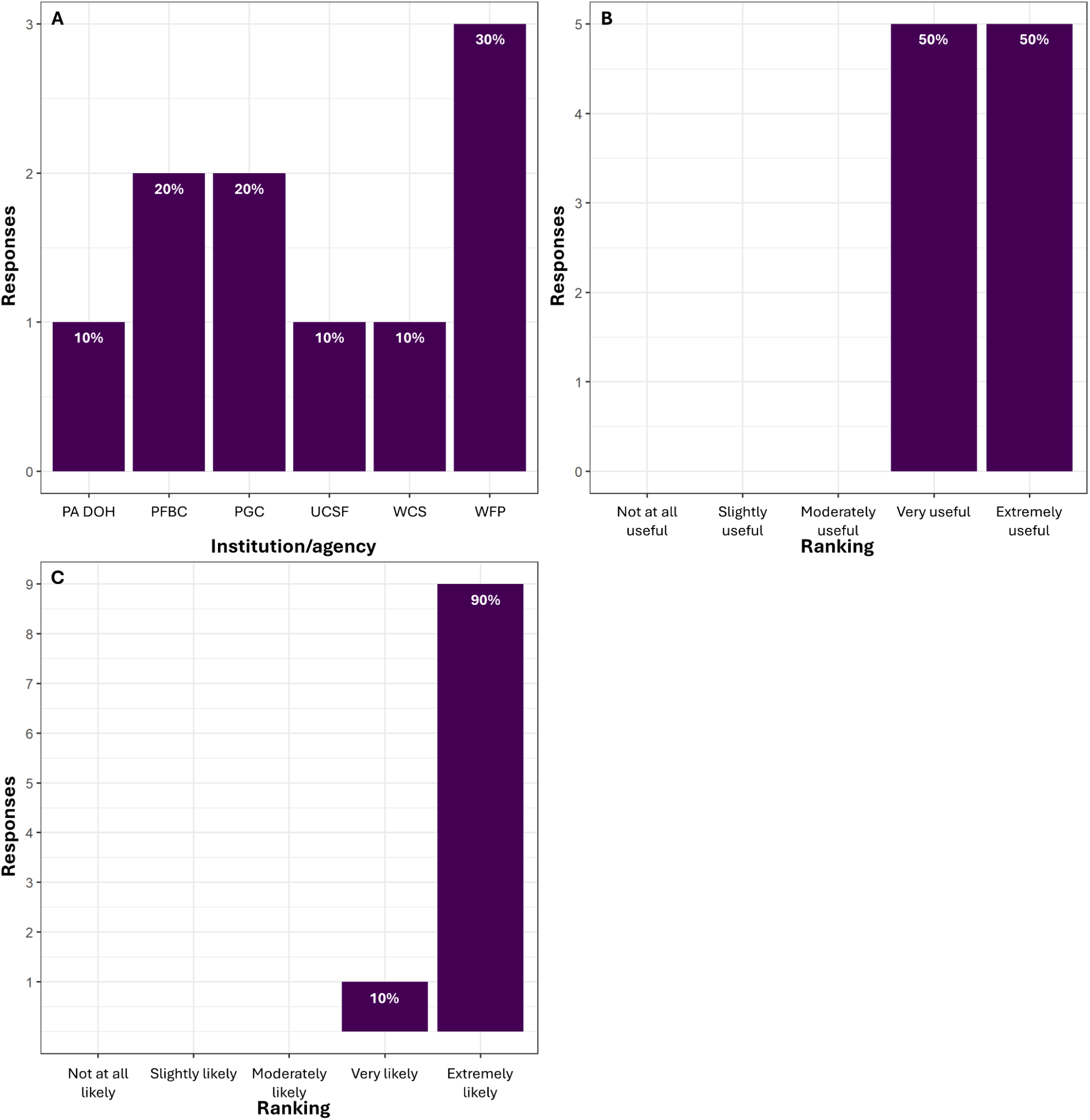
Participant responses (n = 10) to three questions from the tabletop evaluation survey asking A) Which institution or agency they are associated with? B) How useful do they feel the tabletop exercise was? And C) How likely would they be to participate in similar exercises in the future? PA DOH: Pennsylvania Department of Health, PFBC: Pennsylvania Fish and Boat Commission, PGC: Pennsylvania Game Commission, UCSF: University of California San Francisco, WCS: Wildlife Conservation Society, WFP: Wildlife Futures Program.

A clear theme of data sharing and collaboration emerged as a primary takeaway from the event and was highlighted by eight respondents to that survey question. One respondent wrote the biggest takeaway was “That we all need to do a better job, despite all our regulatory limitations and varying agency priorities, of collaborating, at least in a general sense, on how we tackle interagency communication.” Organizers identified three clear themes from the table-top: 1) data sharing between agencies, 2) collective resourcing (e.g., coordinating and sharing personnel and supplies) and 3) and cross-agency relationship building. Respondents were equally split (five each) between ranking data sharing and relationship building as the most important in responding to a disease event, with only a single respondent ranking resourcing as the top priority (Figure 2A). Several respondents noted that relationship building facilitated data sharing, with one respondent noting “All are important. Relationship building is key for facilitating data and resource sharing.”

**Figure 2.**
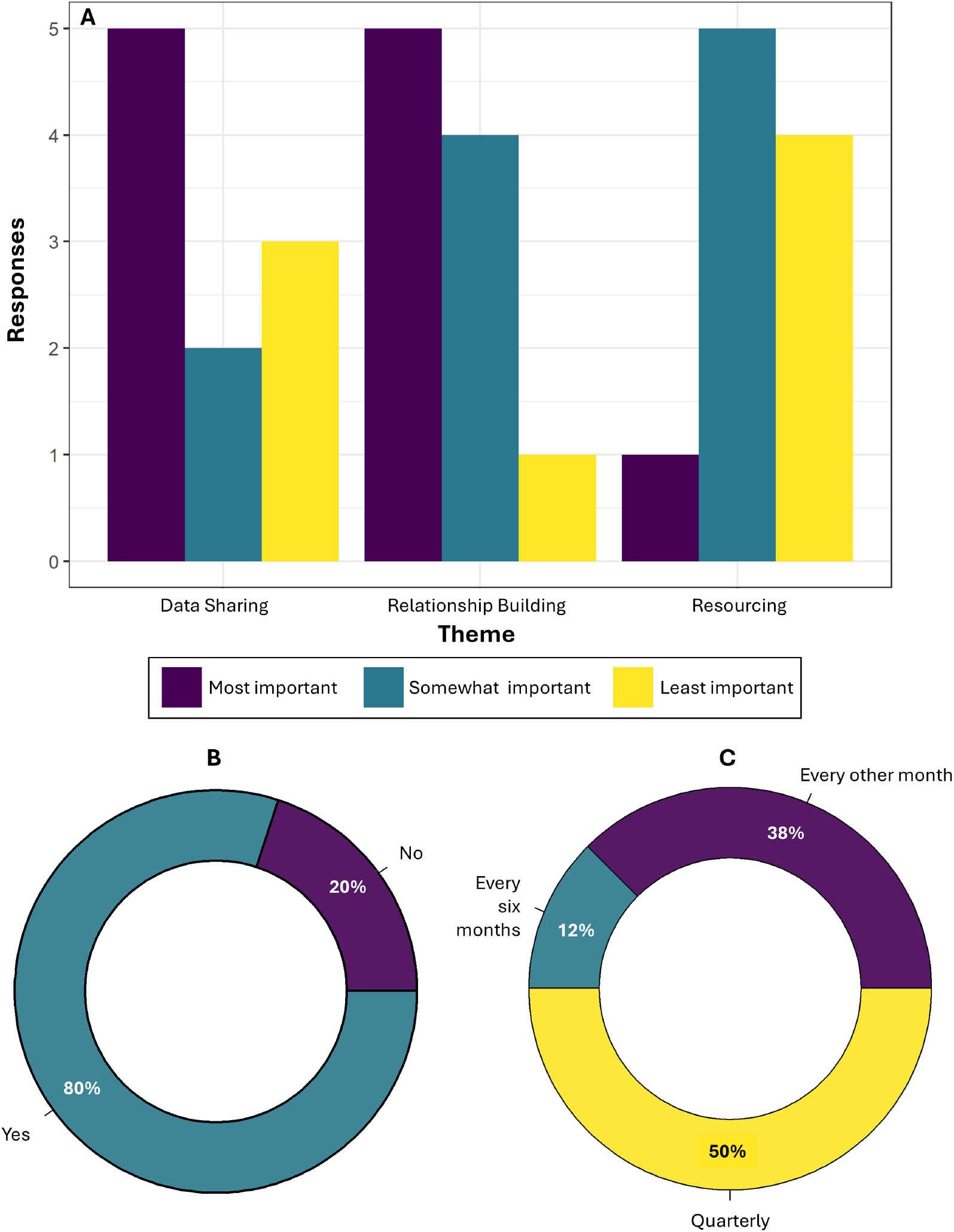
Participant responses (n = 10) to three questions from the tabletop evaluation survey asking A) How would they rank three themes (data sharing across agencies, resourcing e.g., coordination and sharing of personnel, supplies, diagnostic capability, etc., and relationship building e.g., local, interstate agencies, etc.) in order of importance for their role in responding to disease events. B) Would they be interested in joining a working group to bolster interagency preparedness and response to disease events impacting wildlife in Pennsylvania? And C) If they are interested in joining a working group, how often would they be able to participate?

Given the themes that were identified above, participants were asked if they would be interested in joining a working group to improve interagency collaboration for preparedness and response to wildlife disease events. Eighty percent of the respondents said they would join a working group that met regularly, with 50% agreeing that meeting on a quarterly basis would be a good cadence (Figure 2B and C). An additional benefit of a working group is the ability to plan more agency-wide participation in future exercises. One respondent highlighted the potential to actively test this:

> “Would like to see a future exercise actually test our system: we call the field staff and say if this were happening today, how quickly could you respond? What are you capable of doing today? If you’re tied up, who would you call? And then call the lab and say we want to submit xyz, are you able to accept it today? How long of a turn-around would you have, etc.? The group discussion in this exercise was good, especially as an introduction to the players. The next step (next exercise) might benefit everyone by breaking into our own groups and “calling” one another to ask for help, consult, request resources, share information, etc. and then regroup at the end to see how we think it went.”

## Discussion

This exercise brought several challenges to light. Foremost, public-private cooperation is still limited, despite successful examples like the Wildlife Futures Program. Lines of communication exist, but public agencies have substantive limitations on what they are able to financially support, making more formal collaborations and initiatives elusive. Building relationships and trust with state agencies, as well as the private sector, is paramount to strengthening the overall emergency response network. Stronger relationships will also promote collective problem solving around the three priority areas that surfaced during the table-top exercise.

1. Formal collaborative agreements should be put in place through memorandums of understanding before an emergency to enhance cross-agency coordination during an outbreak. These agreements could include SOPs for triggering alert systems and for conducting event and indication-based surveillance. Testing and response protocols during an emergency could be prioritized, with considerations for what types of data each organization needs for decision making and how these might be shared.
2. Proactive designation of approval processes for the mobilizing resources for surge capacity and a chain of command for decision making will reduce friction between competing mandates that may arise during a disease outbreak. Such provisions will also facilitate more efficient cross-organizational collaboration and pooling of resources for effective implementation of mitigation strategies. Additionally, this will help highlight partners with complementary expertise that can be engaged during an emergency and identify areas of redundancy for more efficient use of resources.
3. Fitting each organization’s emergency response plan within the larger inter-agency communication network can build resilience and sustainability by enhancing information flow and data sharing, and reducing reliance on personal relationships to identify points of contact. A lack of communication can lead to missed opportunities for early identification of a potential threat, a delayed response to containment, and public misinformation. Therefore, it is important to first understand what is currently feasible around data sharing and what would be required to improve communication. Making access to data less cumbersome with guidelines for the types of data and when it would be shared will also be a key step to take. Finally, understanding if there are information systems that were set up during the COVID-19 pandemic that could be leveraged now would help facilitate progress.

### Call to action

It is time to fortify our local and state-wide pandemic preparedness and response planning by bringing together a One Health working group to collectively find solutions to common barriers to effective and efficient emergency response. The proposed working group should engage with the Pennsylvania Emergency Management Agency, as well as federal agencies such as the Center for Disease Control and Prevention, for preparedness planning and would perform a variety of key activities that may include generating standard operating procedures for surveillance, putting formal cross-organization collaboration agreements in place, establishing guidelines for communication and data sharing, and generating a pool of emergency response resources that are accessible to the One Health partners. Once infrastructure is established, multi-sectoral collaboration can continue to be reinforced outside of periods of crisis with cross-training exercises, ongoing evaluation and prioritization of research areas and pursuit of funding opportunities.

## Appendix 1

Survey Questions

### Contact information

1. Institution or agency

### Tabletop exercise

2. Briefly state the main reason for your participation in the tabletop exercise.
3. Overall, do you feel the tabletop exercise was useful? For this question, 1 is not at all useful and 5 is extremely useful.
  i. Please elaborate on your rating.
4. What was your biggest takeaway from the day?
5. Three main themes around interagency coordination were identified throughout the day. Please rank these in order of importance for your role in responding to disease events. 1 being most important and 3 being least important.
  i. Data Sharing (across agencies)
  ii. Resourcing (e.g., coordination and sharing of personnel, supplies, diagnostic capability, etc.)
  iii. Relationship Building (local, interstate agencies, etc.)
6. Please elaborate on your ranking.

### Future exercises

7. How likely would you be to participate in similar exercises? For this question, 1 is not at all likely and 5 is extremely likely.
8. What improvements would you suggest for a future tabletop exercise?
9. What other agencies or stakeholders should be invited to a future tabletop exercise? Please list them below.
10. What topics would you like to see future tabletop exercises address?

### Final thoughts

11. Would you be interested in joining a working group to bolster interagency preparedness and response to disease events impacting wildlife in PA?
  A. Yes
  B. No
  C. I don’t know
12. If you are interested, how often would you be able to participate?
  A. Every other month
  B. Quarterly
  C. Every six months
  D. I am not interested
  E. Other
13. Are there any other comments you would like to share?

